# Olfactory-Trigeminal Integration in the Primary Olfactory Cortex

**DOI:** 10.1101/2020.06.24.168989

**Authors:** Prasanna R. Karunanayaka, Jiaming Lu, Rommy Elyan, Qing X. Yang, K. Sathian

## Abstract

Humans naturally integrate signals from the olfactory and intranasal trigeminal systems. A tight interplay has been demonstrated between these two systems, and yet the underlying neural circuitry that mediates olfactory-trigeminal integration remains poorly understood. Using functional magnetic resonance imaging (fMRI), combined with psychophysics, this study investigated the neural mechanisms underlying olfactory-trigeminal integration. Fifteen participants with normal olfactory function performed a localization task with air-puff stimuli, phenylethyl alcohol (PEA; rose odor), or a combination thereof while being scanned. The ability to localize PEA to either nostril was at chance. Yet, its presence significantly improved the localization accuracy of weak, but not strong, air-puffs, relative to air-puff localization without concomitant PEA when both stimuli were delivered concurrently to the same nostril, but not when different nostrils received the two stimuli. This enhancement in localization accuracy, exemplifying the principles of spatial coincidence and inverse effectiveness in multisensory integration, was associated with multisensory integrative activity in the primary olfactory (POC), orbitofrontal (OFC), superior temporal (STC), inferior parietal (IPC) and cingulate cortices, and in the cerebellum. Multisensory enhancement in most of these regions, except the OFC, correlated with behavioral multisensory enhancement, as did increases in connectivity between some of these regions. We interpret these findings as indicating that the POC is part of a distributed brain network mediating integration between the olfactory and trigeminal systems.

**HIGHLIGHTS:**

- Psychophysical and neuroimaging study of olfactory-mechanosensory (OM) integration
- Behavior, cortical activity and network connectivity show OM integration
- OM integration obeys principles of inverse effectiveness and spatial coincidence
- Behavioral and neural measures of OM integration are correlated

## 1. INTRODUCTION

The olfactory and intranasal trigeminal systems convey complementary information that is merged via mutually enhancing and/or suppressive interactions [1, 2]. For example, the trigeminal component of an odorant (at least at higher concentrations) reduces human perception of odor intensities and qualities [3–5]. Conversely, trigeminal stimuli are perceived as more intense in the presence of olfactory co-stimulation [6, 7]. Thus, perceptual judgements in either olfactory or trigeminal modalities partly reflect the influences of other systems. These observations support the premise that the olfactory and intranasal trigeminal systems work in tandem [2].

The ophthalmic and maxillary branches of the trigeminal (fifth cranial) nerve innervate the nasal cavity [8]. Touch, temperature, and nociception are mediated by trigeminal afferents, with different sensory receptors mediating different sensations [9, 10]. Although the mechanical, thermal, and chemical somatosensory domains are processed via the same trigeminal nerve, the corresponding central sensory pathways differ [11]. Olfactory information is independently processed via the first cranial (olfactory) nerve. Since most odorants stimulate both cranial nerves I and V, mutual interactivity between these two systems takes place across multiple processing levels [1, 2, 12].

Multisensory integration improves the detection, discrimination, and categorization of stimuli, conferring evolutionary advantages to animals and humans in understanding, navigating, and perceiving their environments [13, 14]. In the traditional hierarchical view, integration is deferred until *after* extensive ‘unisensory’ processing [15, 16]. The superior temporal sulcus (STS) [17], lateral occipitotemporal [18], inferior parietal (IPC) [19], and ventrolateral prefrontal cortices [20] have all been identified as brain regions where multisensory information converges. Interestingly, recent results suggest a different perspective on multisensory integration, wherein this process is supported by a distributed neural network that includes the primary sensory cortices, in addition to higher-order multisensory brain regions [21, 22].

The olfactory system is often considered an underdetermined human sense [23, 24]. Multisensory integration, therefore, can impart substantial benefits to olfactory processing [25], e.g., the absence of pertinent visual information is known to impede odor identification [24]. Similarly, adding a pure trigeminal stimulus to varying concentrations of an odorant modulates the perceived intensity of the odor [6]. Furthermore, trigeminal perception is significantly impaired in participants with anosmia [26]. These observations point to a tight coupling or mutual reinforcement between olfactory and trigeminal systems, and yet the mechanisms underlying these interactions remain largely unknown [27–31].

The intranasal trigeminal system has been investigated using localization tasks in which participants are asked to identify the stimulated nostril [32–34]. These tasks capitalize on the fact that an odorant can *only* be localized to a nostril if that odorant also stimulates the trigeminal nerve [32]. Thus, localization tasks are ideal for dissociating between the perceptual contributions of trigeminal versus olfactory stimuli. These studies have shown that humans can localize trigeminal and bimodal (olfactory and trigeminal) stimuli, but they cannot localize pure odorants that *only* activate the olfactory nerve [32, 35–37].

Olfactory processing incorporates both sniffing and smelling [38, 39]. Sniffing is typically active, but passive sniffing is a useful laboratory task that allows precise stimulus control and avoids confounds of enhanced odorant delivery due to increased airflow. Both active and passive sniffing improve olfactory perception [35]. During active sniffing, the olfactory system estimates airflow rates in both nostrils using “efference copy” information about the motor action of sniffing and somatosensory input taken from changes in incoming airflow rates [38]. Sobel et al. (1998) reported sniff-induced activation of the primary olfactory cortex (POC) as a result of changes in active sniffing behavior, as opposed to the more passive act of odorant detection. During passive sniffing, an odorant is delivered into participants’ nostrils using air-puffs. Thus, air-puff induced stimulation is closely related to the somatosensory component of active sniffing behavior, which is considered an integral part of the olfactory percept [38, 39]. Although less studied, olfactory information has been shown to influence an intranasal trigeminal percept; olfactory co-stimulation in the same nostril has been shown to enhance the localizability of intranasal trigeminal stimuli for both chemosensory [28] and mechanosensory stimulation [40]. To our knowledge, the effects of such olfactory co-stimulation on the neural processing of intranasal trigeminal stimuli have not been studied.

During passive sniffing, the intranasal trigeminal system is mechanically stimulated by airflow changes. Here we used a psychophysical approach in combination with functional magnetic resonance imaging (fMRI) to investigate the effects of a pure odorant, phenyl ethyl alcohol (PEA; rose odor), which cannot itself be localized to either nostril above chance, on the localizability of intranasal somatosensory stimuli (air-puffs). We hypothesized that multisensory integration would occur for the combination of tactile mechanosensory and olfactory stimuli. Accordingly, we predicted that PEA would enhance localization of weak, but not strong air-puffs, conforming to the principle of inverse effectiveness in multisensory integration, i.e., integration is disproportionately larger for weaker vs stronger crossmodal stimuli [14]. Furthermore, we predicted that PEA would enhance the localizability of weak air-puffs only when both types of stimuli were presented to the same nostril, and not when PEA and air-puffs were presented to opposite nostrils, in keeping with the spatial principle of multisensory integration [41], i.e., integration occurs for spatially concordant but not discordant stimuli. Earlier, we reported psychophysical findings consistent with these predictions, but in the absence of a direct manipulation of air-puff intensity [40].

In the present study, psychophysical data were acquired during fMRI which was used to examine the neural bases of these multisensory interactions. Data from rodents [22], non-human primates [42], and humans [24] suggest that crossmodal stimulation is capable of modulating activity in the POC. This brain structure receives extensive reciprocal projections from other olfactory and higher-order brain regions, such as the orbitofrontal cortex (OFC)—a critical site for multisensory integration from olfactory afferents [43]. We therefore tested the hypothesis that olfactory-trigeminal integration is supported by multisensory interaction in the POC (as in other primary sensory cortices) [15]. It has been suggested that sensory convergence in primary sensory cortices may be mediated by higher-order multisensory brain regions [44]. Hence, we also tested the hypothesis that multisensory effects in the OFC, STS and IPC (known higher-order multisensory brain regions) contribute to olfactory-trigeminal integration. Lastly, we tested the hypothesis that multisensory interaction in the POC is associated with changes in its connectivity with the OFC, STS, and IPC.

## 2. METHODS

### 2. 1. Participants

Fifteen participants (mean age: 27.20 ± 5.07 years, 11 females and 4 males) took part in the fMRI and associated psychophysical experiments. Their smell function was evaluated using the University of Pennsylvania Smell Identification Test (UPSIT) [45] and the OLFACT [46, 47]. UPSIT tests an individual’s ability to detect and identify 40 odorants utilizing “scratch and sniff” strips; participants are given four choices from which to select the name of each odorant. OLFACT is a test battery that evaluates odor identification and odor threshold abilities using a computer-based system. The study had prior approval from the Pennsylvania State University College of Medicine Institutional Review Board (IRB). Participants (recruited from south-central Pennsylvania) provided informed consent and were screened to rule out: (i) MRI incompatibility, (ii) any neurological or psychiatric conditions, and (iii) disorders that could specifically cause olfactory dysfunction, such as viral infections and allergies.

### 2. 2. Imaging Protocol

Weak and strong air-puffs were used in two fMRI runs each. Functional images of the entire brain were acquired using gradient-echo echoplanar imaging on a Siemens Prisma 3.0 T system with a 20-channel head coil. The following imaging parameters were used: repetition time/echo time/flip angle (TR/TE/FA) = 2,000 ms/30 ms/90°; field of view (FOV) = 220 mm x 220 mm; acquisition matrix = 80 x 80; 34 slices; slice thickness = 4 mm, and the number of repetitions = 240. The total scan time of the functional runs was 32 mins (8 min per run). For anatomical overlay, T1-weighted images with 1 mm isotropic resolution were acquired with the magnetization-prepared, rapid gradient echo (MPRAGE) method: TE = 2.98 ms; TR = 2300 ms; inversion time = 900 ms; FA = 9°; FOV = 256 mm × 256 mm × 160 mm; acquisition matrix = 256 × 256 × 160; acceleration factor = 2.

### 2.3. Experimental design

The structure of the paradigm and event timings are shown in **Figure 1**. There were four runs: two runs with weak air-puffs (airflow of 2L/min), and the other two with strong air-puffs (airflow of 4L/min). Run order was pseudorandomized and counterbalanced across participants. For each run, there were four experimental conditions presented pseudorandomly: 8 trials for each condition—totaling 32 trials per run, i.e. 128 trials across all four runs, evenly distributed amongst four conditions. In Condition 1, air-puffs were presented alone while in Condition 2, PEA (2-phenylethyl alcohol; Sigma # 77861) was presented alone, without air-puffs. In each of these two conditions, monorhinal stimuli alternated randomly between nostrils and participants were asked to localize the stimulated nostril via left or right keypress in a two-alternative forced-choice (2AFC) paradigm. Air-puffs were embedded in a constant flow of odorless air, delivered bilaterally at a rate of 2 L/min or 4 L/min, via an olfactometer.

**Figure 1.**
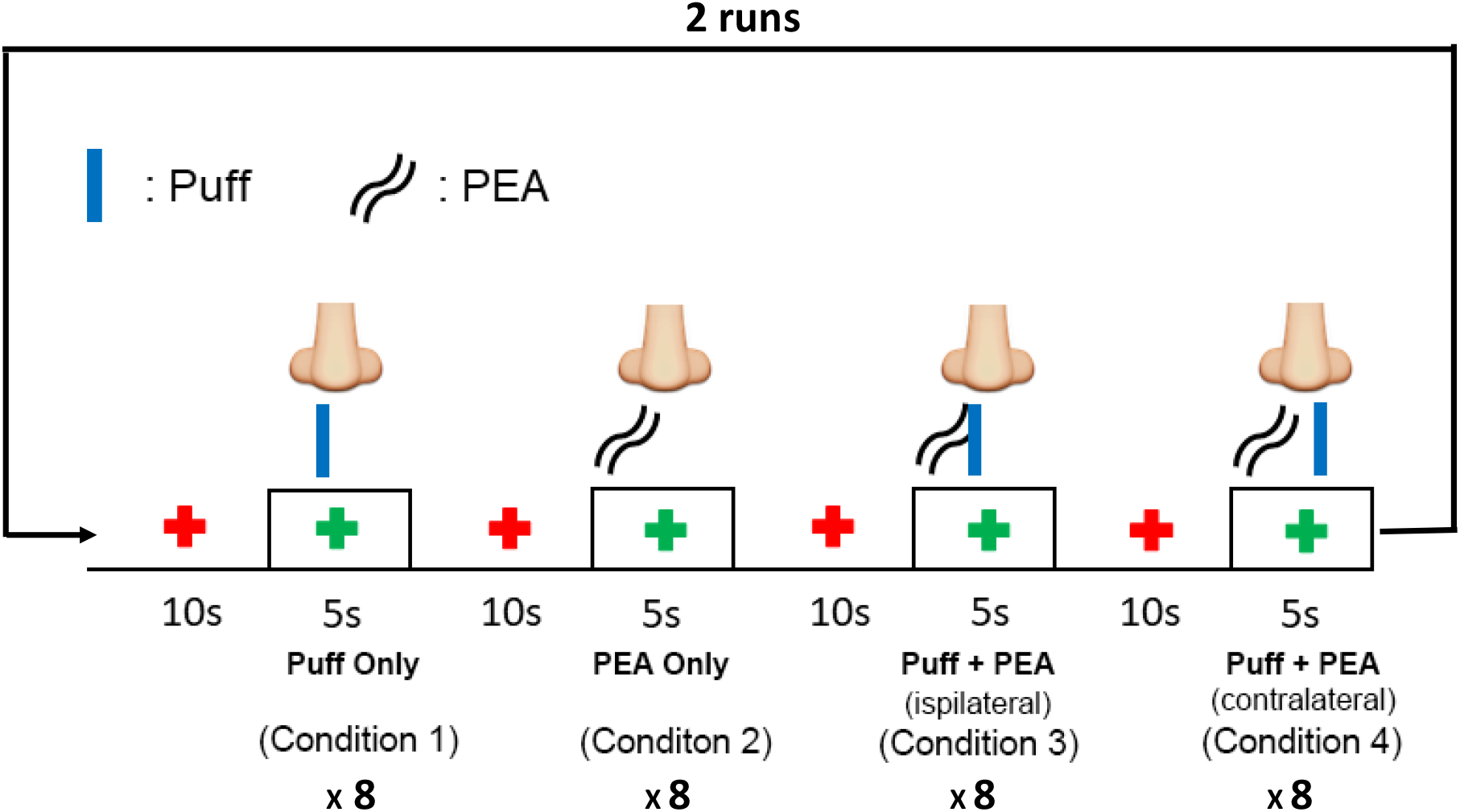
Air-puffs with a flow rate of 2L/min(weak) or 4L/min(strong), and/or pure odorant stimuli (PEA) were randomly delivered to either the left or right nostril, with an inter-stimulus interval of 10 s. Participants were instructed to focus on a red ‘fixation cross’ that appeared on screen. When the cross turned green, they were instructed to hold their breath for 5 s and attend to the stimulus, which lasted 100ms. Participants provided left or right button responses indicating which nostril was stimulated. Responses were digitally recorded with the olfactometer. The paradigm structure was used for weak and strong air-puff runs.

Conditions 3 and 4 were comprised of bisensory stimulation with both air-puffs *and* PEA. In Condition 3, air-puffs and PEA were both presented to the same nostril (ipsilateral), whereas in Condition 4, air-puffs and PEA were presented to opposite nostrils (contralateral). Stimuli alternated randomly between nostrils in conditions 3 (air puff+PEA to same nostril) and 4 (air puff to the left nostril and PEA to the right, or vice versa). In all conditions, participants performed the 2AFC task for localizing the nostril they perceived to be stimulated. The stimulus duration was always 100 ms embedded within a 5 second “breath-holding” period (green cross), while an additional 10 second interval (red cross) was interposed between the 5 s breath-holding periods (see **Figure 1**). Visual cues were used to inform participants to hold their breath, and to localize the stimulated nostril. We carefully selected the inter-stimulus interval (ISI) and the total experiment time to avoid desensitization during the testing period [48]. Five-to-ten-minute pauses between runs were incorporated to ensure that participants were not tired and that their sensory perception remained stable.

### 2.4. Data Analysis

#### 2.4.1. Behavioral Data Analysis

Statistical analyses of behavioral data acquired during imaging were performed using SPSS (IBM SPSS Statistics). We first performed one-sample t-tests to determine whether the localization accuracy for each condition was greater than chance. We then used paired t-tests for comparisons between the various conditions, with Bonferroni corrections for multiple comparisons (corrected alpha value of 0.05).

#### 2.4.2. fMRI Data Processing

fMRI data was processed using SPM 12 (https://www.fil.ion.ucl.ac.uk/spm/software/spm12/). Images were realigned within each run; they were slice-timing corrected and spatially normalized to the Montreal Neurological Institute’s (MNI) EPI brain template [49, 50]. These normalized images were smoothed with an 8×8×8 mm^3^ (full width at half maximum, FWHM) 3D-Gaussian smoothing kernel. Statistical parametric maps were then generated for each subject, and for each condition, by fitting the paradigm time course, convolved with the canonical hemodynamic response function (HRF) and its time derivatives, to the imaging data. One-sample t-tests were conducted to generate group activation maps. Following Beauchamp et al. (2005), multisensory interactions were investigated using a balanced contrast that required the multisensory response to be larger than the mean of the unisensory responses. In SPM, it was implemented using the contrast [−1, −1, 2] (air-puff, PEA, combination). Statistical inferences were performed at the voxel level with an FDR (False Discovery Rate) correction for multiple comparisons (p < 0.05; [51]) across the whole brain, using AFNI software (https://afni.nimh.nih.gov/).

#### 2.4.3. Brain connectivity analysis

The generalized form of the psycho-physiological interaction (gPPI) model is routinely used to examine changes in functional connectivity during specific contexts as defined by the various conditions within the cognitive task [52]. Here, we used gPPI to separately test for functional connectivity changes between brain regions that potentially mediate multisensory enhancement [53], by comparing the unisensory air-puff condition with the bisensory combination of air-puffs and PEA. This analysis was confined to the runs with weak air-puffs, since it was these runs that demonstrated multisensory enhancement (as defined above). For each analysis, the seed region was defined by the group-level activation maps, with the threshold set to p<0.05, FDR-corrected. For each seed region analysis, we first specified PPI models at the subject level using normalized, smoothed, functional EPI images as the data; condition-specific onset times for each run as the ‘psychological’ factor, and the first eigenvariate of seed region activation as the ‘physiological’ factor. These models also included nuisance regressors for the run-wise, motion-correction parameters. Estimated connectivity parameters were entered into group level, one-sample t-tests.

## 3. RESULTS

### 3.1. Behavioral findings

All study participants were normosmic (**Table 1**).

**Table 1.** Olfactory test scores of the study population

|  | OLFACT |  | UPSIT |
| --- | --- | --- | --- |
|  | *Threshold* | *Identification* | *Identification* |
| Mean ± SD | 11.73 ± 1.56 | 17.73 ± 1.33 | 36.27 ± 1.75 |
| Normal Range | $\geq 8$ | $\geq 16$ (Total 20) | $\geq 34$ (Total 40) |

**Figure 2** shows localization accuracy for weak air-puffs alone, which was comparable to that of PEA alone, being below well 75% correct (usually taken as the threshold in a 2AFC). There was a significant improvement in localization accuracy to around 85% correct for weak air-puffs during PEA co-stimulation. However, this improvement was only observed when both stimuli were delivered to the same nostril. No such improvements in localization accuracy were observed when weak air-puffs and PEA were concurrently delivered to *opposite* nostrils. These results suggest that the olfactory-trigeminal interaction observed here fits with the spatial principle of multisensory integration. Further, they rule out the possibility that enhancement of mechanosensory perception by the odorant arose from a non-specific, e.g. alerting, effect. In contrast to the findings with weak air-puffs, strong air-puffs, which could be localized with suprathreshold accuracy in the absence of PEA, showed no significant improvement in localization accuracy when in the presence of PEA (**Figure 2B**), whether PEA was presented to the same or opposite nostril. Of note, localization accuracy for strong air-puffs without PEA was well below ceiling, indicating that the lack of significant improvement with PEA was probably not due to a ceiling effect. These results also suggest that olfactory-trigeminal interactions conform to the principle of inverse effectiveness, i.e., weak stimuli benefit more from multisensory integration than strong stimuli [54–56]. These in-scanner results are similar to those we previously reported in an out-of-scanner behavioral study, although the intensity of air-puff stimulation was not manipulated in the prior study [40].

**Figure 2.**
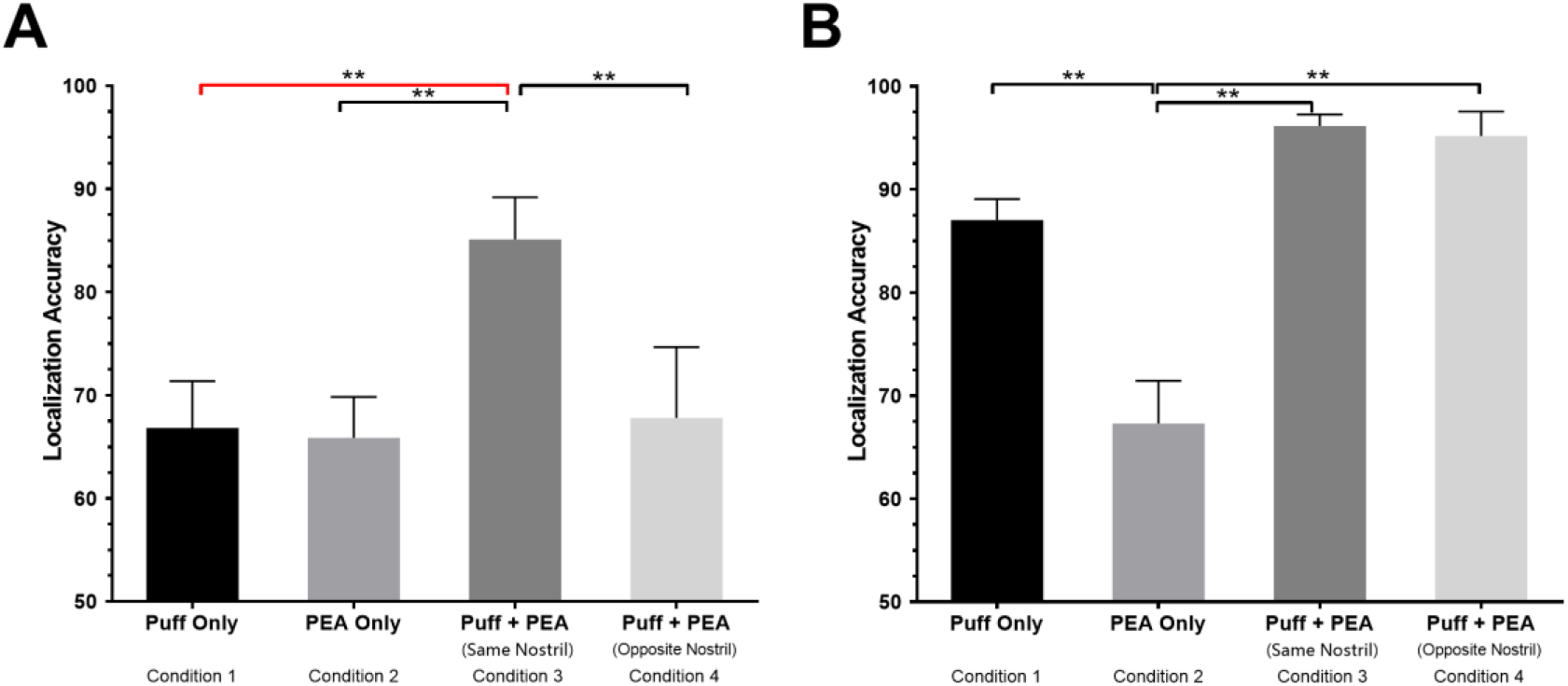
**(A)** Localization accuracy for weak air-puffs significantly improved when PEA was presented to the same nostril (paired t-test, **p < 0.001), but not the opposite nostril. **(B)** Strong air-puffs were localized far better than weak air-puffs, but no significant improvement was observed with PEA co-stimulation. This was the case, whether PEA was in the same or opposite nostrils. Taken together, these results support the **principle of inverse effectiveness** during olfactory-trigeminal integration [38].

### 3.2. Multisensory effects during olfactory-trigeminal interaction

In order to investigate the loci of multisensory integration, we first analyzed fMRI data from the weak air-puff section of the experiment, since these data clearly demonstrate behavioral multisensory integration, embodying both the spatial principle and the principle of inverse effectiveness. Previous studies have suggested that merging olfactory and trigeminal information could be mediated via integrative mechanisms involving several brain regions. It should be noted that classical superadditivity in neurophysiological studies requires that the multisensory response be larger than the summed unisensory responses. However, Beauchamp et al. (2005) showed that this criterion failed to classify the STS as a multisensory region in fMRI studies, though there is an abundance of evidence suggesting that this region integrates multisensory information [57]. Using modified criteria, that the multisensory response should be larger than the maximum or the mean of unisensory responses, STS was classified as multisensory. Following this empirical demonstration [58], we opted to use the criterion of multisensory responses being larger than the mean of unisensory responses to delineate multisensory effects during olfactory-trigeminal interaction. As noted in the Methods section, this was implemented using a balanced univariate contrast [−1, −1, 2] (puff, PEA, combination), with correction for multiple comparisons across the brain. As shown in **Figure 3,** several loci satisfied this criterion for olfactory-trigeminal integration. These loci were in parts of the primary olfactory (piriform), superior temporal, orbitofrontal, inferior parietal, and cingulate cortices, as well as in the cerebellum. The activations were bilateral, except for the superior temporal and orbitofrontal activations, which were on the left. Critically, such multisensory integration was only found for weak air-puff stimuli in the presence of PEA; strong air-puffs evoked responses that were not modulated by the addition of concomitant PEA. Additionally, multisensory enhancement was only present when PEA and weak air-puffs were presented to the same nostrils, being absent when PEA and weak air-puffs were presented to opposite nostrils. This conformance to the principles of inverse effectiveness and spatial congruence matches the corresponding behavioral effects and reinforces the idea that the observed fMRI activity patterns reflect multisensory integration of olfactory and intranasal mechanosensory inputs.

**Figure 3.**
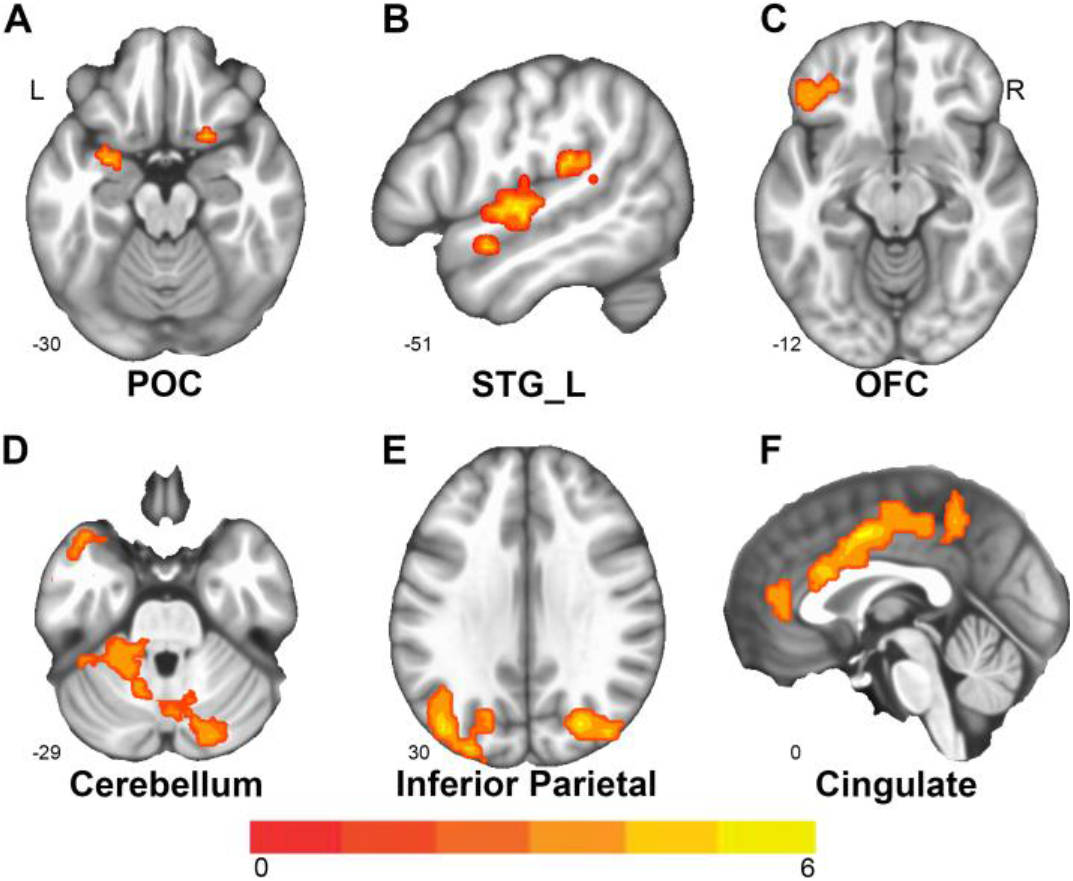
Multisensory effects in the POC (A), left superior temporal cortex (B), left OFC (C), cerebellum (D), IPC (E),and cingulate cortex (F) (p<0.05, FDR corrected) during the weak air-puff experiment. No such multisensory effects were observed for the strong air-puff experiment (data not shown).

### 3.3. Multisensory effects in the POC during olfactory-trigeminal interaction

It is particularly interesting that we identified significant multisensory effects in the piriform cortex, suggesting that this structure may be involved in merging olfactory and trigeminal information even though it is regarded as the primary olfactory cortex. As shown in **Figure 4**, the mean beta (β) value in the POC satisfied our selected criterion for multisensory integration while also conforming to the inverse effectiveness principle.

**Figure 4.**
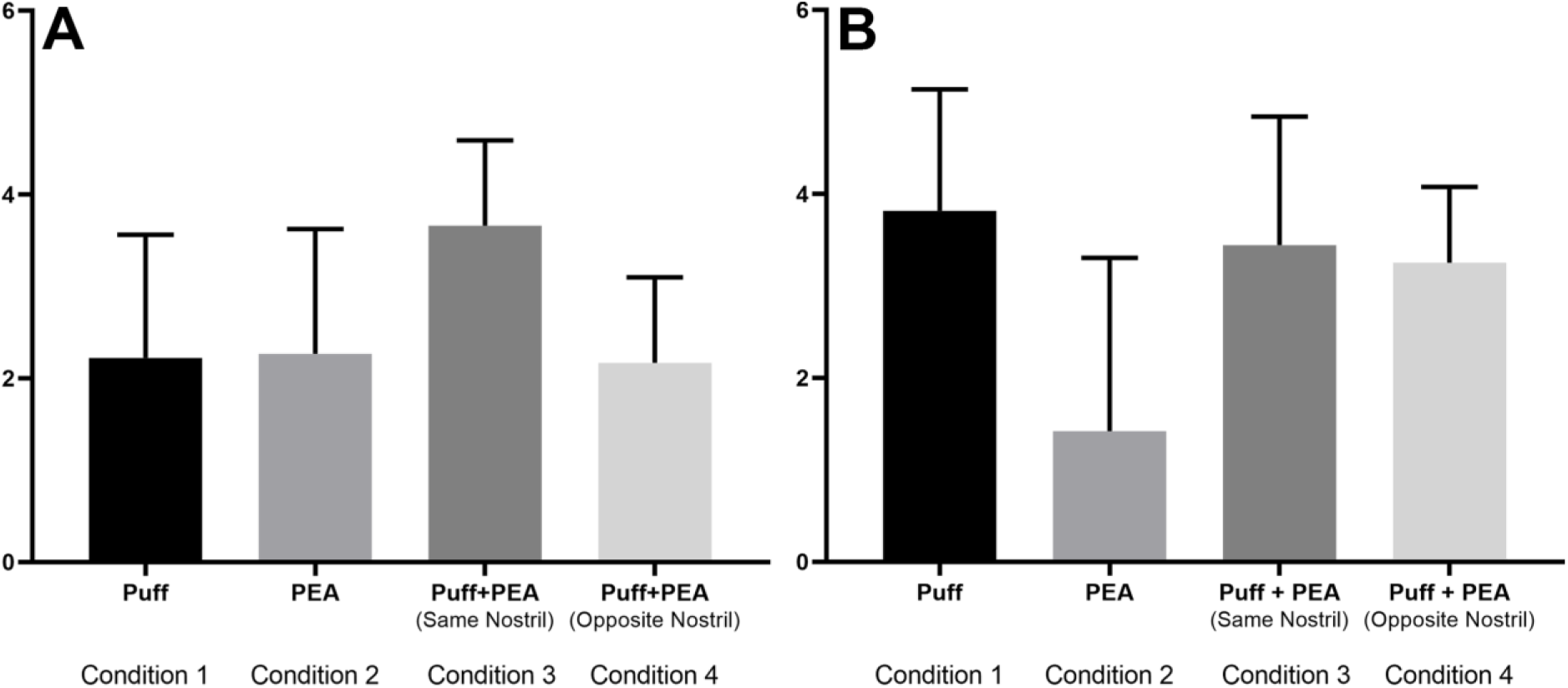
POC response in terms of beta (β) values during air-puff, PEA and air-puff+PEA conditions. **(A)** Weak air-puff experiment showing multisensory enhancement. **(B)** Strong air-puff experiment showing no multisensory enhancement.

We further analyzed the β value distribution within the activated POC volume. **Figure 5** shows that certain voxels (red) satisfy the commonly used criterion for “superadditivity”, i.e., multisensory responses are larger than the sum of the unisensory responses [58]. Our data suggests that multisensory effects in the POC may be heterogeneous in nature, and that some voxels exhibit stronger multisensory effects than others.

**Figure 5.**
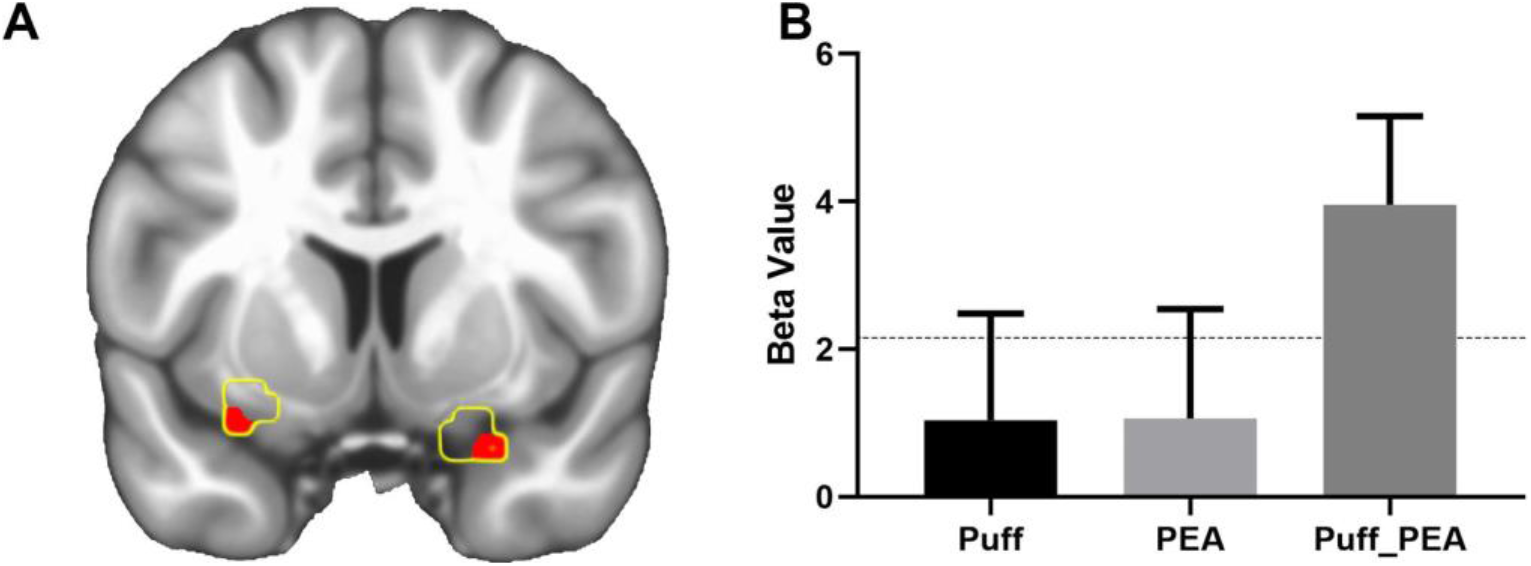
Parameter estimates (β) within the POC (A) during olfactory-trigeminal integration. POC activity in response to weak air-puffs or PEA (B), and their combination, in a subregion of the POC (red).

### 3.4. Multisensory effects in other brain regions during olfactory-trigeminal interaction

We also identified multisensory effects in the superior temporal cortex and the IPC (**Figure 3**). The STS/STG is the auditory association cortex, and has been implicated in binding auditory, visual, and somatosensory information [57]. This region could serve a similar role in merging olfactory and trigeminal information [59]. The IPC’s role as a general filter and integrator of sensory information, within and across modalities, is well recognized. The inferior parietal lobule and intraparietal sulcus (IPS) could potentially subserve attention for a specific modality over others [60]. For example, in visual, auditory, and tactile modalities, the IPS appears to facilitate processes such as crossmodal localization and spatial attention [16, 61]. We suggest a similar role for the IPC during our localization experiments. As seen in **Figure 3**, we also observed multisensory effects during olfactory-trigeminal integration in the cerebellum. Previously, this structure has been implicated in olfactory processing. The observed cerebellar activity in this study may be related to a feedback mechanism linking and modulating intranasal somatosensory and odor information [62].

### 3.5. Behavioral correlations of fMRI multisensory effects

While the present findings provide evidence for multisensory effects at the behavioral and neural levels, it remains unclear whether multisensory response enhancement in the POC and other areas, relative to unisensory responses, actually predicted the extent of multisensory behavioral enhancement. To address this issue, we performed correlation analyses by regressing subject-specific multisensory fMRI enhancement (i.e., air-puff+PEA combination minus the mean of unisensory responses) against multisensory enhancement of changes in localization accuracy (for weak air-puffs+PEA combination minus air-puff alone). There was a significant correlation between improvement in localization accuracy and the magnitude of multisensory enhancement in the left POC, left superior temporal cortex (STC), left IPC, mid-cingulate cortex (MCC), and left cerebellum (**Figure 6**). No such correlation was observed between the magnitudes of multisensory enhancement in OFC and localization accuracy. These findings support the idea that multisensory effects in the POC, STC, IPC, MCC, and cerebellum may underlie the observed behavioral enhancements during the weak air-puff localization experiment.

**Figure 6.**
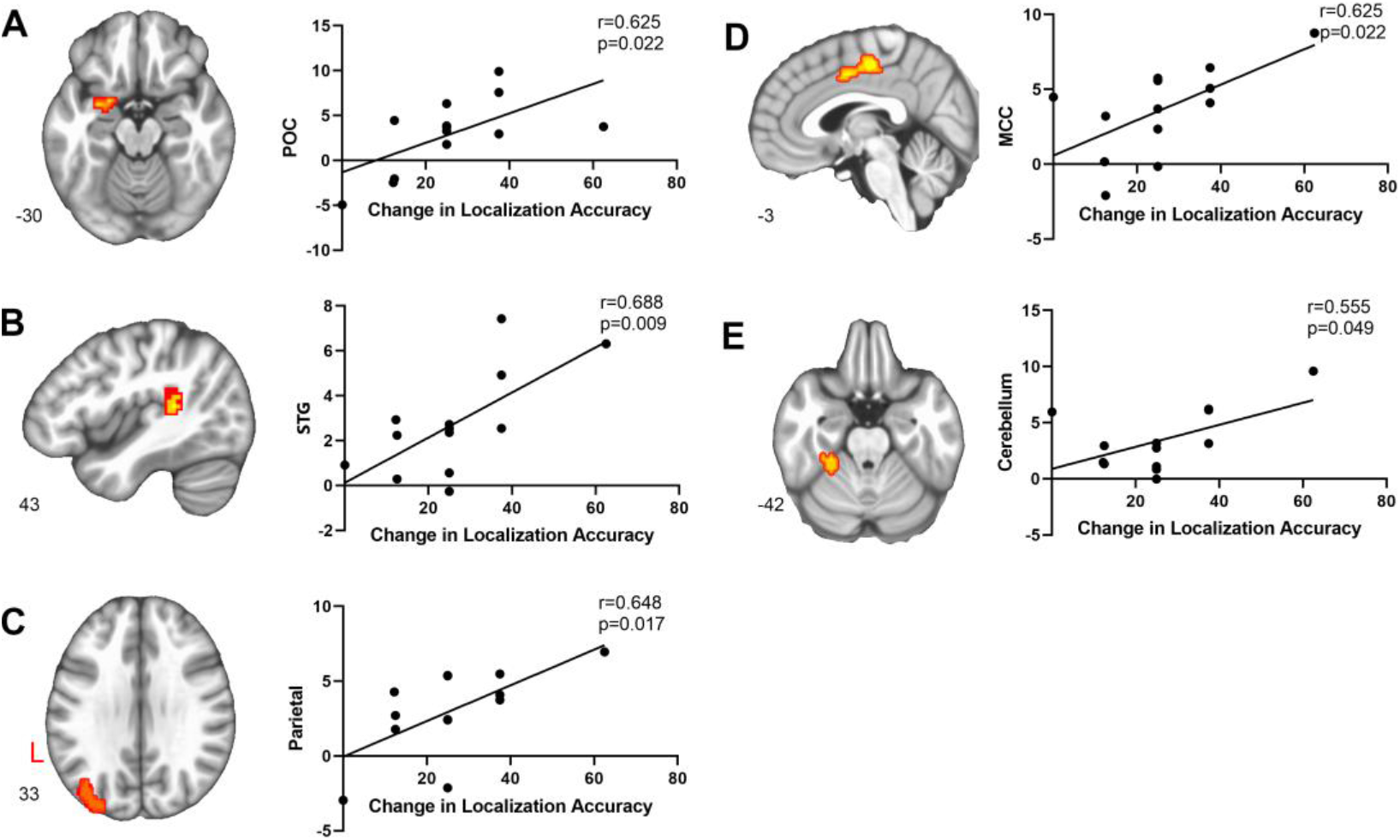
Brain-behavior relationships. Scatterplots demonstrating strong correlations between multisensory fMRI activity and improvements in localization accuracy.

### 3.6. Pathways mediating olfactory-trigeminal interaction

Previously, Gottfried et al. (2003) identified a region of the OFC (adjacent to the POC) that is involved in merging olfactory and visual information. We therefore, considered the possibility that changes in propagation of olfactory information, from the POC to the OFC, might account for the enhanced behavioral effects observed during the weak-air puff experiment. To test this possibility, we implemented connectivity analysis using the generalized psychophysiological toolbox [53]. In this analysis, we tested for multisensory-related modulation of connectivity of the regions demonstrating multisensory enhancement, as shown in **Figure 3**(see Methods). To be precise, we used separate gPPI analyses to compute functional connectivity changes between each of the regions shown in Figure 3 and the whole brain for the air-puff+PEA condition minus the weak air-puff only condition. We then tested how these connectivity changes were related to changes in localization abilities between conditions. Three seed regions (in the POC, STC, and IPC) exhibited significant connectivity changes, as shown in **Figure 7.** POC connectivity with both the OFC and cerebellum were modulated during olfactory-trigeminal integration. Similar connectivity modulations were observed for the STC and IPC, each with the OFC and cerebellum (see **Figure 7**).

**Figure 7.**
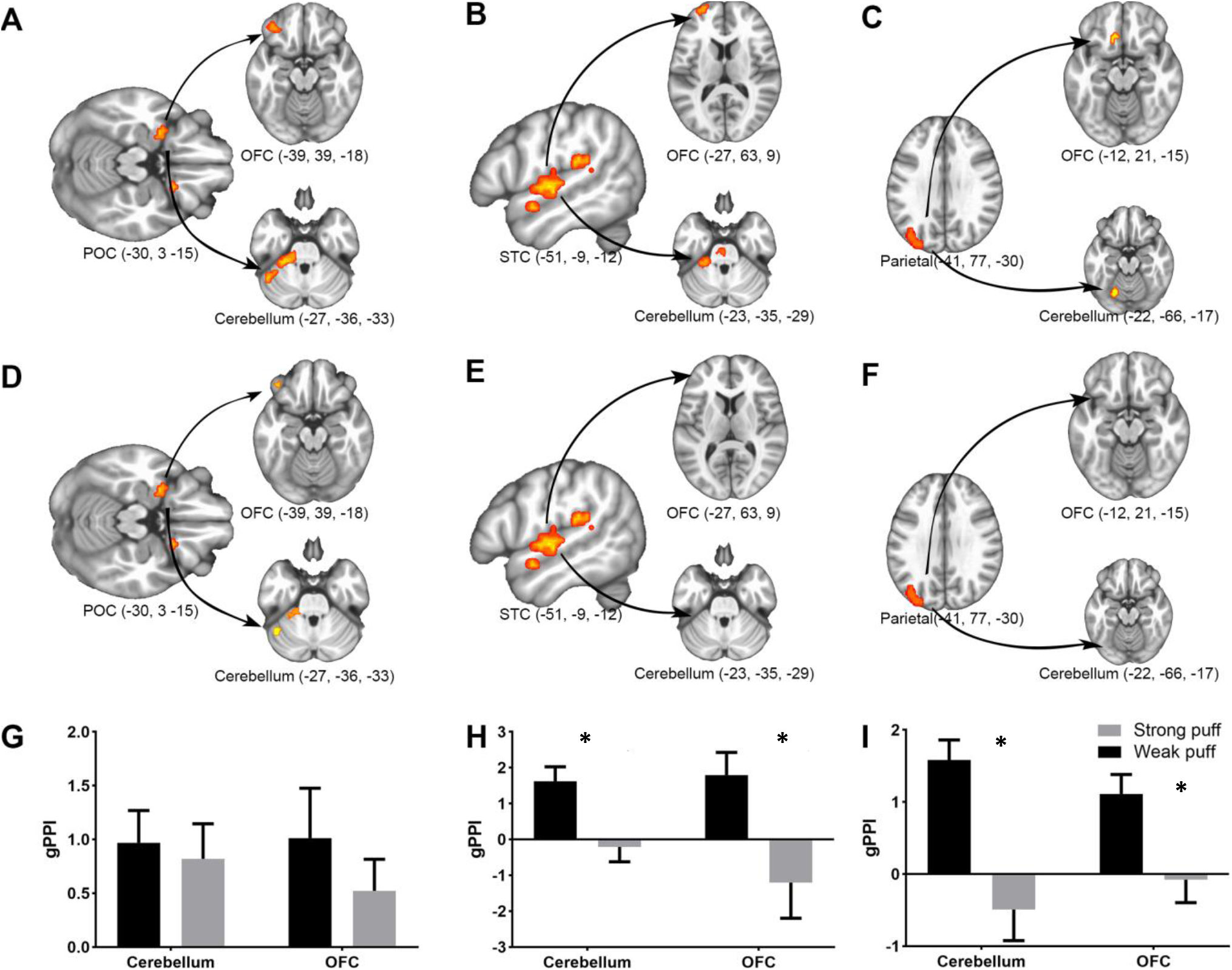
Connectivity changes, i.e., gPPI estimates of the POC, STC, and IPC. These seed regions were selected from clusters of activation showing multisensory enhancement in **Figure 3**. The gPPI connectivity (**A**) from the POC to the OFC and from the POC to the cerebellum; (**B**) from the STC to the OFC and cerebellum; and (**C**) from the IPC to the OFC and cerebellum, during olfactory-trigeminal integration (thresholded at P < 0.05 corrected). The ROIs used in the weak air-puff experiment were used to estimate the gPPI connectivity for the strong air-puff experiment (**D**, **E,** and **F**). gPPI estimates at peak voxels in (**G**) the POC-coupled cerebellum and OFC; (**H**) the STC-coupled cerebellum and OFC; and (**I**) the IPC-coupled cerebellum and OFC. gPPI estimates from the same peak voxels for the strong air-puff experiment are also included. *, P < 0.05 corrected. Note; gPPI is an estimation of functional connectivity differences between two given conditions (e.g., weak air-puff + PEA minus air-puff only).

Interestingly, the multisensory-specific increases in POC–OFC and POC–cerebellar connectivity were positively related to changes in localization accuracy, as shown in **Figure 8**. However, while the change in STC connectivity to the OFC showed no significant correlation with respect to changes in localization accuracy, its connectivity change to the cerebellum showed a negative correlation with changes in localization accuracy. These observations provide support for the hypothesized role of the cerebellum in olfaction: serving as a feedback mechanism that monitors and acquires sensory information for modulating sniffing behavior [62]. Taken together, these findings suggests that functional couplings of the POC and other multisensory areas may support behavioral enhancement during olfactory-trigeminal integration.

**Figure 8.**
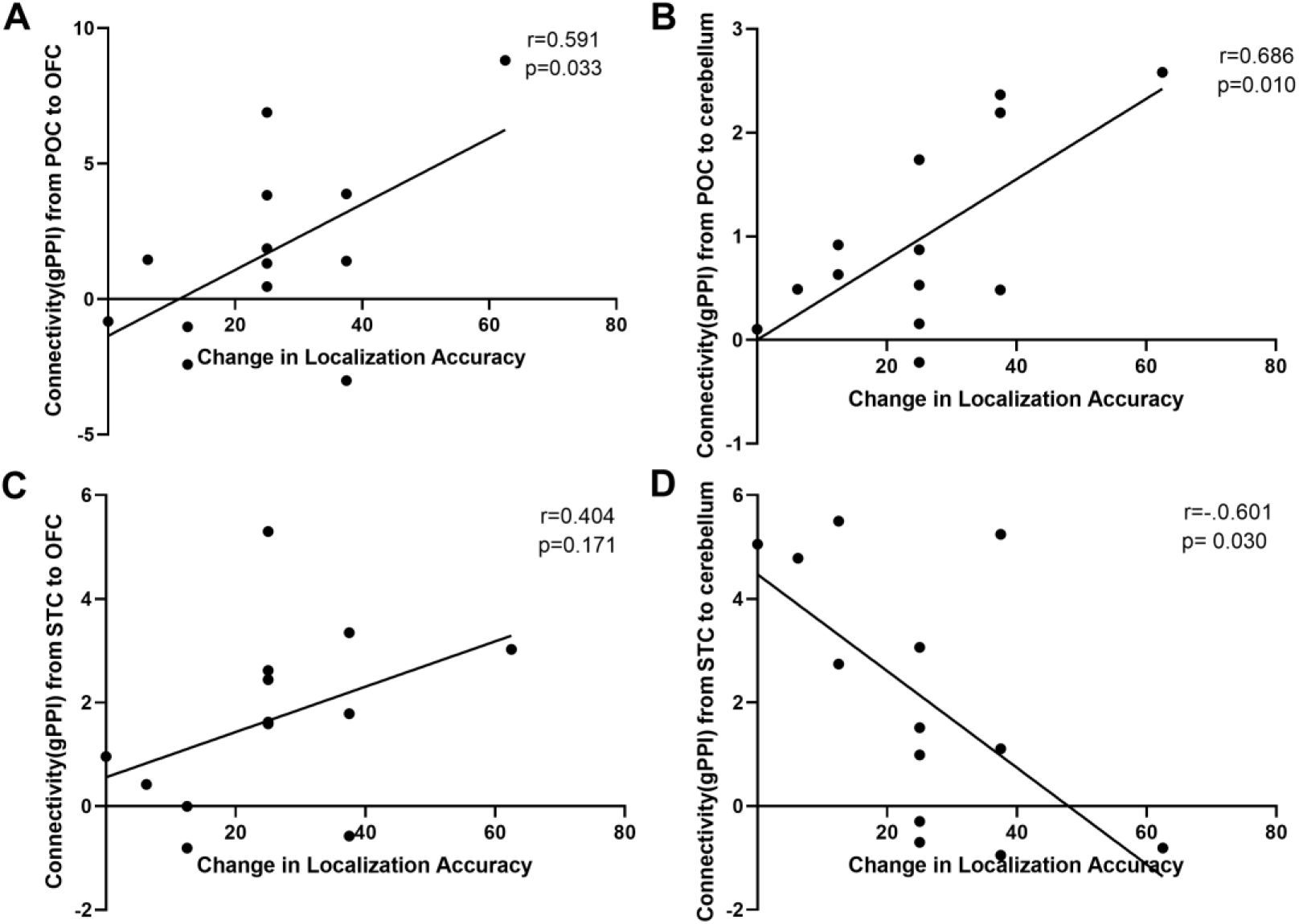
Correlations between gPPI connectivity and behavioral multisensory enhancement. gPPI connectivity between **(A)** the POC and the OFC, and **(B)** the POC and cerebellum were significantly positively correlated with changes in localization accuracy. However, gPPI connectivity estimates between **(C)** the STC and the OFC were not significantly correlated, whereas gPPI estimates between **(D)** the STC and the cerebellum were significantly negatively correlated, with changes in localization accuracy.

## 4. DISCUSSION

### Multisensory integration of olfactory and intranasal mechanosensory information

The ability to merge different sensory information into coherent percepts is a fundamental ability of the central nervous system. This ability is important for the human olfactory system, which strongly relies on binding crossmodal information to form olfactory percepts [63, 64]. Here, we addressed an unresolved question: how does the human brain integrate information from the olfactory and trigeminal systems? We demonstrated olfactory-trigeminal integration behaviorally, using a task calling for localization of intranasal air-puffs to one or other nostril. Although the odorant PEA could not itself be reliably localized, its combination with weak air-puffs significantly enhanced their localizability, but only when the combination was presented to a single nostril and not when PEA and air-puffs were presented to opposite nostrils. Such behavioral enhancement was not found for strong air-puffs. These findings conform to the spatial and inverse effectiveness principles of multisensory integration [14, 41].Our results are also in accord with the temporal principle, given that PEA and air-puffs were presented simultaneously, although we did not test this formally by manipulating synchrony.

fMRI scanning during task performance revealed multisensory integration in the POC; a cortical network comprising the OFC, STC, IPC and MCC; and the cerebellum. Activity patterns in these regions during air-puff localization obeyed the inverse effectiveness principle, since concomitant PEA enhanced the activity evoked by weak, but not strong, air-puffs. They also followed the spatial principle, since enhancement was only found when PEA and air-puff stimuli were presented to the same nostril, and not opposite nostrils. Importantly, the magnitude of multisensory enhancement in all these regions, except the OFC, correlated with the magnitude of behavioral improvement conferred by PEA on the localization of weak air-puffs. Further, connectivity of the POC, STC, and IPC with the OFC and cerebellum exhibited multisensory enhancement. Finally, the magnitude of such multisensory enhancement of POC connectivity with the OFC and cerebellum was positively correlated, and of STC-cerebellar connectivity was negatively correlated, with multisensory improvement of localization accuracy. Our findings suggest that a distributed brain network, which includes the POC, underlies the integration of olfactory and mechanosensory information. This study, therefore, expands our understanding of the neural substrates underlying multisensory integration in an olfactory context.

The trigeminal nerve (cranial nerve V) conveys both chemosensory and somatosensory intranasal trigeminal stimuli, but the associated perceptions appear to be relatively independent, behaviorally [35]. The quality of odors is independently conveyed via the olfactory nerve (cranial nerve I), whose inputs are modulated in the presence of trigeminal co-stimulation [3, 4]. The trigeminal system has been extensively investigated using odor localization tasks, which aim to stimulate the trigeminal nerve [28]. Unlike many behavioral experiments demonstrating response enhancement, our forced-choice localization task with PEA provided an important control for attentional focus, since behavioral and fMRI effects were only present when odorant and mechanosensory stimuli were received by the same nostril. If the effect of adding PEA to weak air-puffs had been obtained even when the stimuli were in opposite nostrils, this could have reflected a non-specific alerting effect. Therefore, the demonstrated effects of multisensory integration were dissociated from the effects of attention. Our results provide compelling evidence for the functional significance of neural responses to combinations of odor and intranasal mechanical stimuli, as well as the existence of central loci where olfactory and trigeminal information are integrated. The impetus for the present fMRI study was provided by our prior behavioral study of olfactory interactions with intranasal somatosensory stimuli [40] and a related behavioral effect of interactivity between olfactory and chemosensory trigeminal stimuli [28]. Future studies should seek to clarify the underlying neural mechanisms of the latter kind of integration.

Passive stimulation by air-puffs closely mimics airflow changes inside the nasal cavity due to sniffing [39]. Sniffing is an active process of stimulus transport, and is considered to be an integral and necessary part of the olfactory percept. For example, odor intensity and identity percepts are modulated by patterns of sniffing [38]. It has been shown that POC activity is driven by the mechanosensory component of sniffing, i.e., airflow changes in the nostrils. The main contribution of our study is that modulation of the perception of somatosensory stimuli, by olfactory information, is associated with neural signals in the POC, related multisensory neocortical areas, and the connections between these areas.

### Multisensory integration in the POC

The complex interplay between the olfactory and intranasal trigeminal systems endows humans with the ability to process information from distinct sensory inputs within the nasal cavity [30]. Based on intensity evaluations, it has been proposed that this interaction may take place centrally [6], although there is support for peripheral interactivity at the mucosal level [30]. Research on multisensory integration in the olfactory system has mostly been conducted in rodents and, to a lesser extent in humans. It has been shown that both gustatory [65] and auditory [22] stimuli can modulate single-unit recordings of neuron activity in primary olfactory areas. Using phase synchrony, Zhou et al. (2019) showed that primary olfactory cortical areas support multisensory integration within the olfactory system [66].

The complex interplay between the olfactory and intranasal trigeminal systems endows humans with the ability to process information from distinct sensory inputs within the nasal cavity [30]. Based on intensity evaluations, it has been proposed that this interaction may take place centrally [6], although there is support for peripheral interactivity at the mucosal level [30]. The present study focused primarily on central aspects of olfactory-trigeminal interaction. The POC has traditionally included brain regions that receive direct afferent inputs from the olfactory bulb, which includes the piriform cortex (PC; a 3-layered paleocortical area), olfactory tubercle, and entorhinal cortex [67, 68]. Our findings challenge the classical dogma that olfactory and trigeminal information must reach known multisensory-specific brain regions *before* being integrated into logical and coherent perceptual experiences, or qualia. Rather, we demonstrate that the POC is itself a locus of multisensory integration, which is consistent with research showing multisensory processing in other primary sensory cortices (e.g., visual, auditory and somatosensory) [13]. This work therefore brings concepts of olfactory processing into line with recent advances in understanding of virtually every sensory system – these advances have led to the idea that multisensory processing is ubiquitous in the brain, at least in the neocortex [15]. We also found that, while the POC as a whole satisfies the criterion that multisensory response magnitude exceeds the mean of unisensory responses, a subset of voxels with the POC exhibit true superadditivity. This suggests that responses within the POC are heterogeneous, which in turn implies that the application of multivoxel pattern analysis methods are likely to be fruitful in further uncovering the function of this region.

### Multisensory integration in the extended olfactory network

A human fMRI study by Gottfried et al. (2003) showed that visual facilitation of olfactory detection enhanced the activity of the OFC, which is distinct from, but interconnected with, the POC [69]. Simultaneous presentation of pure trigeminal and olfactory stimuli led to activations in the OFC and insula, although this study did not demonstrate the integration of olfactory and trigeminal stimuli in the POC [70]. Taste stimulation has also been shown to induce responses in the rodent PC [71], and auditory tones have been known to induce responses in the rodent olfactory tubercle [22]. Although the olfactory bulb (OB) is known to subserve olfactory-trigeminal integration [27, 72–74], its structure cannot easily be imaged with fMRI at 3T. Given the technological limitations, we did not focus on the OB in this study, despite its relevance to olfactory–trigeminal integration.

The POC has reciprocal connections with the OFC, a structure that showed multisensory effects during olfactory-trigeminal integration [75, 76]; visual facilitation of olfactory detection has been shown to enhance activity in heteromodal OFC regions [24]. The OFC is also a critical site for flavor processing because of its sensitivity to olfactory, gustatory, visual, and mouthfeel sensory inputs [77]. As proposed by Kareken et al (2004), OFC activity in the present study may be related to olfactory exploration, or more generally, expectancy [78]. Our study suggested that the POC is a crucial site for olfactory-related associative processing. We then reasoned that multisensory effects in the OFC might be explained by its connectivity with the POC. Indeed, our results supported this hypothesis because the change in connectivity between the POC and OFC reflected localization accuracy of weak air-puff stimuli.

Additional activations, dependent on task performance, were seen in multisensory areas such as the STC, IPC, and MCC. The STS is known to subserve the crossmodal binding of features in auditory and visual modalities. The enhanced STC activation in our study may be related to the crossmodal integration of features between olfactory and trigeminal stimuli, rather than the task of localizing the stimulated nostril. On the other hand, the heteromodal IPC is known to mediate the crossmodal localization of objects in space, which may be relevant to the weak air-puff localization task. Our data shows that response patterns in the STC and IPC are relevant to chemosensation, and that these brain regions (long classified in non-olfactory terms) may in fact be involved in a far wider spectrum of sensory information. Finally, the midcingulate cortex (MCC) contributes to cognitive control, multisensory control, and decision making, but its specific role in olfactory-trigeminal integration remains unclear and needs to be thoroughly investigated in future research [79].

Although activity in the cerebellum during olfactory stimulation has previously been reported, its role in relation to the chemical senses remains unclear [62, 80]. Studies in individuals with olfactory deficits have also highlighted a role for the cerebellum in olfaction (e.g., Bitter et al. 2010a, 2010b; Peng et al. 2013). These findings have led to the premise that (e.g., Bensafi et al. 2008; Iannilli et al. 2011) this structure is part of a complex integrative system (comprising the cerebellum, the entorhinal cortex, sensory areas, and frontal regions), which is related to the intensity coding of olfactory stimuli [81, 82]. Likewise, Small et al. (2003) reported intensity-dependent cerebellar activity (irrespective of valence) during gustatory stimulation, and suggested a role for the cerebellum in processing stimulus intensity during chemosensation. Previous research also suggests a role for the cerebellum in multisensory integration [16, 83–85]. Non-human primate data also suggest that different sensory areas of the cerebral cortex converge upon common areas within the neocerebellum [86–88]. Multisensory activity changes in the cerebellum positively predicted behavioral enhancements, although connectivity changes with the STG negatively predicted behavioral enhancement. The hypothesized involvement of the cerebellum as a feedback mechanism during olfactory processing is consistent with our data [5, 79, 80, 89]. For example, its ability to maintain a feedback loop regulating sniff volume in relation to odor concentration [78]. Our findings imply that the POC, STC, IPC, cerebellum, and MCC are all regions contributing to the formation of the multisensory olfactory percept. Future studies should clarify the functional contributions of these brain structures.

### A neural perspective of olfactory-trigeminal integration

Some olfactory receptors are capable of stimulating olfactory sensory neurons, which makes these neurons responsive to chemical and mechanosensory stimuli [89, 90]. Airflow-driven mechanosensory stimuli are detected by olfactory sensory neurons, which can cause changes in how an olfactory stimulus is percepts [52, 91, 92]. Animal studies have shown that passive sniffing is capable of driving mitral and tufted cell activity in the olfactory bulb, informing downstream olfactory structures about airflow changes in the nasal passages [93–95]. In addition, piriform cortex neurons in mice have demonstrated the capacity to differentially encode olfactory versus trigeminal stimuli [96, 97]. A recent human psychophysical study by Yao et al. (2021) showed that central olfactory processing does incorporate contralateral nasal airflow information; this was hypothesized to be relayed via branches of the trigeminal nerves [98]. Noting that the bilateral olfactory tracts and piriform cortices are interconnected via the anterior olfactory nucleus, the findings by Yao et al. may be explained by the inter-hemispheric transfer of olfactory and/or mechanosensory information, whereby contralateral nasal airflow may have impacted olfactory perception [96, 99–101].

While their study demonstrates that contralateral trigeminal information affects olfactory processing, the converse has yet to be investigated to its fullest extent. Our study showed that contralateral olfactory information did not necessarily add or facilitate the processing of intranasal trigeminal information. In summary, although trigeminal information can inform an olfactory percept regardless of the side of stimulation [as seen in Yao et al. (2021)], our study showed that olfactory stimulation did not aid in the processing and perception of *contralateral* trigeminal (air-puff) stimuli.

### Clinical implications and conclusions

Participants with olfactory deficits often exhibit decreased trigeminal sensitivity [102]. Sinceolfactory information can influence the perception of air-puff stimuli, we hypothesize that somatosensory trigeminal sensitivity may be affected in anosmic patients [103]—challenging the notion that chemosensory and somatosensory systems are relatively independent. Our results, and those of Tremblay et al. (2019), support a general mechanism by which the olfactory system interacts with the intranasal trigeminal system. The POC is a potential brain structure for this interaction and integration, regardless of whether the olfactory information is being merged with chemosensory or somatosensory trigeminal information. As a result, we predict that multisensory effects in the POC might be reduced for both chemosensory and somatosensory stimulation in anosmic participants. Such findings would support the hypothesis that the trigeminal system’s responses are both independent of which trigeminal receptors are being stimulated, or of the type of stimulation [104]. If so, then measuring the sensitivity of one trigeminal stimulus may suffice as a general assessment of the trigeminal system as a whole [36].

Apart from our main objective: determining the multisensory integrative circuitry for olfactory and trigeminal systems, this study provides a basis by which several critical barriers to progress in the field may be addressed. (1) Aging affects the interdependencies of the olfactory and intranasal trigeminal systems; thus, understanding integration deficiencies may be a crucial first step in delineating the aging effects in both systems [105]. (2) Olfactory deficits are preclinical symptoms in Alzheimer’s disease (AD) and Parkinson’s disease (PD) [106–109]. The mechanisms of multisensory integration (between olfactory and trigeminal systems) might be differentially affected in AD and PD. Such information would be of great clinical and research interest: it may be crucial for developing biomarkers capable of identifying and monitoring people at risk for developing AD or PD.

Our current results have identified a distributed activity pattern across a multisensory network that may collectively support olfactory-trigeminal integration. These results also indicate that the basic principles of multisensory integration are preserved across brain structures and modalities. This data highlights the importance of the POC in integrating olfactory and intranasal trigeminal stimuli—reemphasizing its role as an associative cortex [68, 90]. Such early integration allows decision-making to be based on multisensory information, with an initiation of responses faster than would otherwise be possible [14]. The results seem to suggest that (as is the case with vision, audition, gustation, and olfaction) spatial coincidence is a key factor in determining whether integration is taking place between the olfactory and trigeminal systems. Moreover, such integration conforms to the principle of inverse effectiveness. It is also likely to conform to the principle of temporal coincidence, although this remains to be investigated. We found that the integration of olfactory and intranasal mechanosensory information relies on enhanced activity in the POC, in addition to other known higher-order multisensory brain regions. Connectivity between these regions seems to play a fundamental role in integrating information between the two systems.

## ACKNOWLEDGEMENTS

The study was supported by the Leader Family Foundation, a grant from the U.S. National Institute of Aging (R01-AG027771) and the Department of Radiology, Penn State College of Medicine. This project also received partial funding from a grant with the Pennsylvania Department of Health using Tobacco CURE Funds. We would also like to thank Q.X. Yang for their help in revising and editing the manuscript.

## AUTHOR CONTRIBUTIONS

Conceptualization, P.K. and K.S; Methodology, J.L, Q.X.Y. P.K., and K.S.; Investigation, J.L, R.E., P.K., and K.S.; Writing–Original Draft, J.L, R.E., P.K., and K.S.; Funding Acquisition, Q.X.Y. and P.K.; Resources, Q.X.Y. and P.K.; Supervision, P.K. and S.K.

## DECLARATION OF INTERESTS

The authors report no competing interests. The Pennsylvania Department of Health specifically disclaims responsibility for any analyses, interpretations or conclusions.

## SUMMARY

Olfactory and intranasal trigeminal systems influence each other’s performance via suppressing and enhancing interactions. Karunanayaka et al. show that behaviorally relevant multisensory enhancement, in a set of cortical regions including primary olfactory cortex (POC), emerges when humans attempt to localize weak air-puffs in the presence of phenylethyl alcohol (rose odor). Connectivity between the POC and higher-order multisensory brain regions seems to play a role in integrating information between olfactory and trigeminal systems. Behavioral and neuroimaging results suggest that the spatial coincidence and inverse effectiveness principle are key factors influencing integration between these two systems. These findings advance our understanding of the neural processes by which olfactory-trigeminal networks integrate information.

## Notes

### Competing Interest Statement

The authors have declared no competing interest.

